# Structural features of mTORC2 that control substrate-specific activities

**DOI:** 10.1101/2022.02.03.478898

**Authors:** Zanlin Yu, Junliang Chen, Enzo Takagi, Feng Wang, Bidisha Saha, Xi Liu, Lydia-Marie Joubert, Catherine E. Gleason, Mingliang Jin, Chengmin Li, Carlos Nowotny, David Agard, Yifan Cheng, David Pearce

**Author notes:** Correspondence (D.P.). Contributed equally (Zanlin Yu, Junliang Chen).

## Abstract

mTORC2 is a multi-subunit kinase complex that is central to multiple essential signaling pathways. Two core subunits, Rictor and mSin1 distinguish it from its relative, mTORC1 and support context-dependent phosphorylation of its substrates. mTORC2 structures have been determined previously, however, important questions remain, particularly regarding structural determinants of substrate specificity and context dependent activities. We used cryo-EM to obtain high resolution structures of the human mTORC2 apo-complex, as well as structures in the presence of substrates, Akt and SGK1. Specific predictions suggested by substrate-induced structural changes were tested in functional assays. First, side chain interactions between Rictor and mTOR that prevent recruitment of mTORC1 substrates and confer resistance to the mTORC1 inhibitor rapamycin were visualized for the first time in the apo-state, demonstrating the steric occlusion that prevents mTORC2 interaction with mTORC1 substrates and rapamycin. Also in the apo-state, mSin1 was seen to form extensive contacts with Rictor, including a pair of short α-helices nestled between two Rictor helical repeat clusters, followed by an extended strand, which makes multiple weak contacts with Rictor helical cluster 1. In co-complex structures, SGK1, but not Akt, markedly altered the conformation of the mSin1 N-terminal extended strand, disrupting multiple weak interactions while inducing a large rotation of mSin1/Arg-83, which comes to interact with a negative patch within Rictor. Mutation of Arg-83 to Ala selectively disrupted mTORC2 dependent phosphorylation of SGK1 but not of Akt, supporting context-dependent substrate selection. These findings provide new structural and functional insights into mTORC2 specificity and context-dependent activities.

## Introduction

Mechanistic target of rapamycin (mTOR) is a serine/threonine kinase which belongs to the phosphatidylinositol 3-kinase (PI3K) related kinase (PIKK) family and is evolutionarily conserved from yeast to human (Heitman et al., 1991; Sabatini et al., 1994). mTOR signaling is central to a large array cellular processes, mediating responses to hormones and growth factors, nutrient availability, and extracellular ionic milieu (Fu and Hall, 2020). mTOR functions in cellular physiology through two structurally and functionally distinct multiprotein complexes, mTOR complex 1 (mTORC1) and mTOR complex 2 (mTORC2) (Gaubitz et al., 2016; Luo et al., 2018; Saxton and Sabatini, 2017). These two protein complexes share two conserved components, mTOR and mammalian lethal with SEC13 protein 8 (mLST8, also known as G*β*L) (Kim et al., 2003), as well as several complex-specific components. mTORC1 distinctively contains regulatory-associated protein of mTOR (Raptor) and proline-rich Akt substrate of 40 kDa (PRAS40) (Kim et al., 2002; Wang et al., 2007), while mTORC2 specifically includes rapamycin-insensitive companion of mTOR (Rictor), mammalian stress-activated protein kinase-interacting protein 1 (mSin1). These compositional differences underlie the distinct substrate preferences of the two complexes, as well as their differential responses to the macrolide immunosuppressant, rapamycin; mTORC1 is acutely and potently inhibited by rapamycin while mTORC2 responds only partially after long-term treatment (Loewith et al., 2002; Sarbassov et al., 2004; Sarbassov et al., 2006).

The function of mTORC2 has received increasing attention during recent years. The representative downstream effectors of mTORC2 are mainly members of the AGC kinase family, including Akt (also known as protein kinase B), protein kinase C (PKC), and serum- and glucocorticoid-induced kinases (SGK1 and SGK3) (Cameron et al., 2011b; Li and Gao, 2014; Lu et al., 2010; Malik et al., 2018; Wu et al., 2011). Upon activation by mTORC2, these AGC kinases have been demonstrated to play fundamental roles in regulating cell metabolism, proliferation, survival, migration, and ion transport (Albert et al., 2016; Chellappa et al., 2019; Gleason et al., 2015; Kazyken et al., 2019; Sato et al., 2016; Sorensen et al., 2019). Dysregulation of mTORC2 signaling has been reported in some human diseases such as cancer, metabolic disorders, and neurodegenerative diseases (Johnson et al., 2015; Srivastava et al., 2019; Tang et al., 2016; Venugopal et al., 2020; Yuan et al., 2017).

In part because it was discovered first and in part due to its inhibition by rapamycin, functional and structural research on mTORC1 has outpaced mTORC2. Cryo-EM and X-ray crystallographic structural studies of mTORC1 have substantially elucidated its structural features, in most domains at the atomic level. Important structural determinants of substrate-specificity and rapamycin sensitivity have been elucidated. Notably, a domain within mTOR close to the catalytic cleft, termed the FKBP12-Rapamycin (FRB) domain, binds both rapamycin (in complex with the endogenous FK-506 binding protein, FKBP12) and key mTORC1 substrates, 4EBP1 and S6-kinase (Yang et al., 2013).

mTORC2 structures have been determined by cryo-EM as well, but multiple key structural features, most notably those that confer context-dependent substrate phosphorylation remain to be elucidated. Comparison of published structures indicates that the common subunits, mTOR and mLST8, assume very similar conformations in both mTORC1 and mTORC2, suggesting that specificity determinants lie elsewhere. Chen et al. used cross-linking mass spectrometry coupled with a 4.9 Å cryo-EM structure to identify interactions between the mTOR FRB domain and mTORC2 components, mSin1 and Rictor (Chen et al., 2018). Their data suggested that both of these mTORC2-specific subunits closely interacted with the FRB domain in a manner that could block binding of FKBP12-Rapamycin, as well as that of the substrates, 4EBP1 and S6-kinase. However, structural detail was lacking. Scaiola et al. (Scaiola et al., 2020), published a higher resolution structure which provided greater insight into the FRB domain, as well as clarifying the position and interactions of mSin1 with Rictor and mLST8. Furthermore, they showed that mSin1 did not interact directly with mTOR, but rather provided an important link between Rictor and mLST8. However, significant detail of the structure of the mSin1 N-terminal region was not visualized. This is particularly important in the context of recruitment of specific substrates. Prior functional and biochemical studies had identified two distinct substrate interaction domains in mSin1: one in the N-terminal region of mSin1 was found to be specific for SGK1 and did not interact with Akt or PKC (Lu et al., 2011b). Mutation of residues within the core of this domain disrupted mTORC2-dependent phosphorylation of SGK1 but not Akt or PKC. A second domain, the CRIM (conserved region in the middle) domain, was found to be required for binding and activation of all three of these substrates (Cameron et al., 2011b). The role of the CRIM domain was further elucidated in an NMR-based study (Tatebe et al., 2017). However, the N-terminal region has not been further characterized. Structural insights into the N-terminal region and CRIM domain have been elusive.

In this study, we present a representative cryo-EM structure of human mTORC2 apo-complex at overall resolution of 3.2Å together with structures in the presence of Akt and SGK1. We describe the characteristic assembly that provides a basis for modelling local structural information including recognition of side chains in multiple regions. The structure reveals the basis for mTORC2 insensitivity to rapamycin, demonstrating specific side chain interactions through which Rictor occludes the mTOR FRB domain. Structural determinants of substrate preferences of mTORC1 and mTORC2 are also revealed. Of particular note, new insight into the structural basis for mTORC2 context-dependent regulation of SGK1 and Akt is revealed: in the presence of SGK1, but not Akt, the conformation of the mSin1 N-terminal region is altered, resulting in a large rotation of mSin1/Arg-83 to form a salt bridge with Rictor Asp-1679. The functional importance of this newly recognized structural feature is demonstrated in cell-based phosphorylation experiments.

## Results

In order to obtain purified mTORC2 core complex for cryo-EM, Spycatcher003-mTOR, Flag-Rictor, HA-mLST8 and mSin1-HA were co-expressed in Expi293F™ cells. The complex was purified using one-step flag pull-down, followed by size exclusion chromatography (Figure S 1A, B and C), and finally subjected to on-grid affinity binding using a new adaptation of the Spycatcher-Spytag system (Wang et al., 2020), before cryo-EM analysis. This combined approach allowed high quality cryoEM samples to be obtained without needing large-scale expression and purification approaches. Our principal sample was prepared in the apo-state without small molecule additives or peptide substrates. The biochemical activities of purified mTORC2 were tested and verified (Figure S1D). After 2D and 3D data analysis, approximately 200,000 particles were selected and used for the final refinement and reconstruction (Figure S2). Based on the density map data, a model was built for the mTORC2 apo-state (Figure S1E). Similar structures were solved for co-complexes of mTORC2 with two of its principal substrates, Akt and SGK1.

### Global aspects of mTORC2 structure

The mTORC2 structure comprises eight subunits consisting of two heterotetramers showing a 2-fold symmetry, similar to a prior report (Chen et al., 2018). The overall shape of the apocomplex is that of a rhomboid with a hollow center and a dimension of 220 × 160 × 140 (Å^3^) (Figure 1A and B). Each of the heterotetramers consists of mTOR, Rictor, mLST8 and mSin1 with a stoichiometry of 1:1:1:1. The two mTOR subunits form a dimer that serves as the foundation of the complex. mLST8 and Rictor sit in close proximity on its two distal ends. The structures of mLST8 and mTOR, as well as their mutual interaction pattern are quite similar to that in mTORC1 (Aylett et al., 2016; Yang et al., 2013). The kinase loop of mTOR is located in the cleft between mLST8 and Rictor. Interestingly, as further addressed below, mSin1 forms a long bridge-like structure connecting mLST8 and Rictor, which has an extended α-helix that crosses over the deep cleft harboring the mTOR active site. 3D variability analysis showed that mTORC2 is not a rigid complex, and the two halves were found to “breathe” in two modes, mainly twisting or squeezing toward the geometric core of the complex (Figure 1B). Both movements result in the altering of the interaction between the two symmetric mTOR subunits, as shown dynamically in supplemental movies (Supp. Movie 1 and 2).

**Figure 1.**
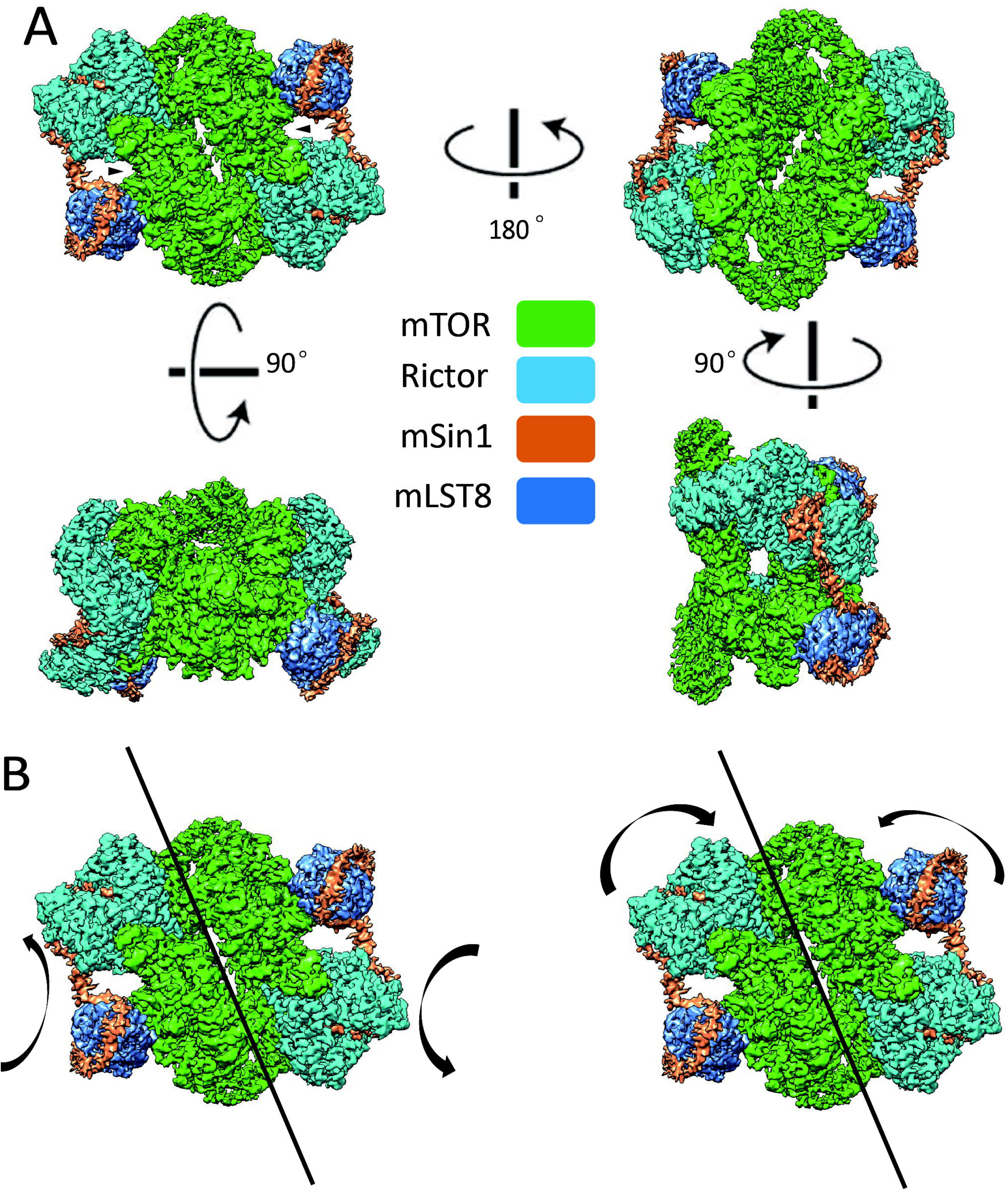
The overall structure of mTORC2: (A) Cryo-EM structure of mTORC2 at overall resolution 3.2Å was reconstructed. The sample was prepared in the apo-state. Four views of the structure are presented, and the rotation axis and degree between views are indicated. The subunits are colored as indicated: mTOR (green), Rictor (cyan), mLST8 (blue), and mSin1 (orange). The kinase loop of mTOR is shown by arrows. (B) Movement of the two heterotetramers of mTORC2 along the central axis are in two directions, twisting (left panel) and squeezing (right panel) toward the geometric core of mTORC2. See also supplementary data for movies.

### Specific side chain interactions underlie Rictor contribution to mTORC2 substrate specificity and resistance to rapamycin

In the apo-complex the two mTOR subunits are anchored to each other, forming the core of the complex and having a conformation quite similar to that in mTORC1 (Figure 1A). The structure of Rictor includes three HEAT (Huntingtin, Elongation factor 3, protein phosphatase 2A, and the yeast kinase TOR) helical repeat (HR) clusters termed HR1, HR2, and HR3 (Figure 2A), all of which make direct contacts with mTOR. HR1 and HR2 also interact with mSin1. Consistent with previous biochemical and lower resolution cryo-EM results, Rictor and mLST8 do not directly interact, but rather are held together by mSin1.

**Figure 2.**
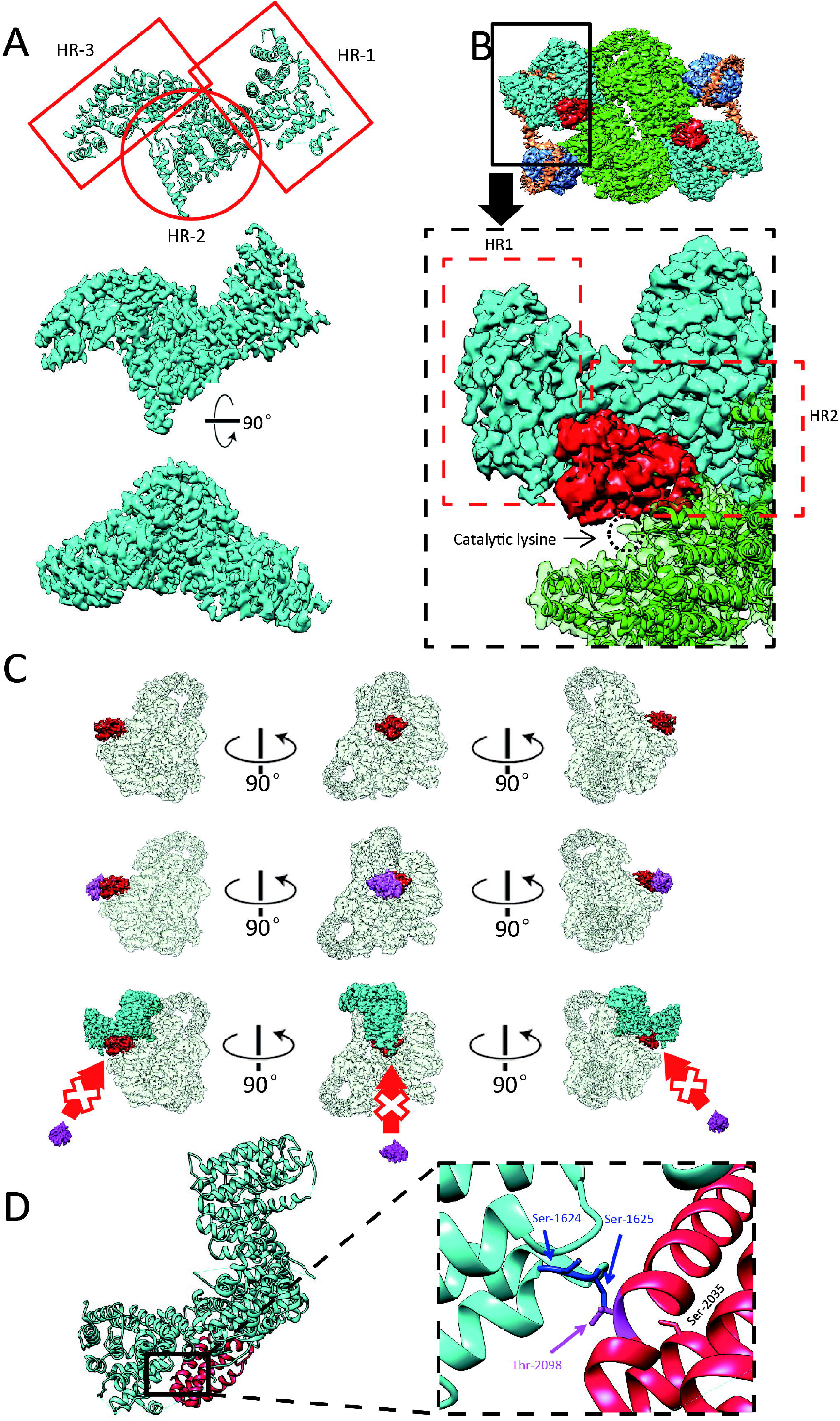
Rictor interaction with mTOR blocks rapamycin accessibility and influences mTORC2 substrate specificity: (A) The atomic model based on density map generated by cryo-EM (upper panel). The high-resolution density map ensures the accuracy of the model fitted from two representative views (mid and lower panel). (B) The cryo-EM density of FRB domain is colored in red in the overall density map (upper panel). In the close-up view, HR1 and HR2 of Rictor are highlighted by red dashed rectangle frames, and the mTOR catalytic lysine is highlighted by a black dashed circle located in the kinase cleft as indicated. To focus on the interaction of Rictor with FRB domain, mSin1 is not shown here. (C) Upper panel: Three views of the density map of mTOR (transparent except FRB domain shown in red). Middle panel: Putative FKBP12-rapamycin binding with FRB domain regardless of Rictor. Lower panel: FRB domain becomes inaccessible to FKBP12-rapamycin in the presence of Rictor. (D) The atomic model of Rictor (cyan) and FRB domain of mTOR (red), as shown in the left panel. The close-up view of the local area illustrates where Thr-2098 on FRB domain and its relevant Ser-1624 and Ser-1625 of Rictor are located (right panel). Ser-2035 is shown as well.

In Figure 2B, the FRB (FKBP-rapamycin complex binding) domain of mTOR is highlighted in red to call attention to its interaction with Rictor. In mTORC1, this domain is adjacent to both the kinase active site and to the RNC (Raptor N-terminal Conserved) domain of Raptor, which plays a central role in substrate recruitment (S6-Kinase and 4EBP (Yang et al., 2013)), and is involved in binding the FKBP12-rapamycin inhibitory complex (Aylett et al., 2016). In mTORC2, the FRB domain is similarly located adjacent to the mTOR kinase active site, but also interacts with both HR1 and HR2 of Rictor. Importantly, HR1 of Rictor sterically prevents the FRB domain from interacting with the FKBP12-rapamycin complex (Figure 2C), while in mTORC1, the FRB domain remains accessible (Figure S6). This steric occlusion also prevents mTORC2 interaction with mTORC1-specific substrates and shielding a key interaction residue, Thr-2098 (Yang et al., 2017). Rictor/Ser-1624 and Ser-1625 were observed in close proximity to mTOR/Thr-2098, likely contributing to the interaction of Rictor HR1 and the FRB domain through hydrogen bond formation, and hence to the differential specificities of mTORC1 and mTORC2 (Figure 2D). Furthermore, mTOR/Ser-2035, which was previously shown to play an important functional role in substrate recognition by mTORC1 (McMahon et al., 2002; Yang et al., 2013), is spatially shielded by Rictor, although there is no direct interaction between them (Figure 2D). Finally, neither SGK1 nor Akt had any observable effect on the relationship of Rictor and mTOR.

### mSin1 engages in key interactions with both Rictor and mLST8

Based on functional and biochemical data, mSin1 contains four principal domains: an N-terminal domain, CRIM domain, RBD (Ras binding domain) and PH (pleckstrin homology) domain (Frias et al., 2006). For the present apo-complex, we solved the major structure of the N-terminal domain, while the other three domains are invisible in the current density map likely due to their flexibility, as has been noted in prior publications (Chen et al., 2018; Scaiola et al., 2020). Interestingly, one of the non-visualized domains, the CRIM domain, has been implicated in binding of all three major mTORC2 substrates, Akt, SGK and PKC. The N-terminal domain (aa 1-137) contains three distinct sections: a Rictor-interacting section, a bridge section connecting Rictor and mLST8, and an elongated section wrapping around mLST8 (Figure 3A). The Rictor-interacting section includes a pair of short α-helices nestled between HR1 and HR2 of Rictor, followed by an extended strand, which runs along the periphery of HR1, making multiple weak contacts (Figure 3A, B). This region, which we have termed the String (SGK-Targeting, Rictor-InteractING) domain has not been visualized in prior density maps, which is notable in light of the role it plays in substrate binding. Gln-68, in particular, was previously shown to be necessary for SGK1—but not Akt or PKC—interaction with and phosphorylation by mTORC2 (Lu et al., 2011a). In the apo-complex, the Gln-68 side chain is well resolved and seen to reach toward Rictor/Arg-105, consistent with a stabilizing interaction (Figure 3C). At the C-terminal end of the string domain, we see Thr-86 embedded in a negatively charged pocket of the Rictor C-terminal region, and specifically interacting with Rictor/Glu-1675. Interestingly, Thr-86 lies within the sequence RRRSNT, which conforms to the consensus substrate peptide for AGC kinase (sequence: RXRXXS/T), and has been shown previously to be phosphorylated by Akt and S6-kinase (Liu et al., 2013), however, in the present structures (including co-complexes), it is unphosphorylated. Downstream of the string domain, an α-helical “bridge” begins immediately C-terminal to Thr-86, and crosses the deep cleft harboring the mTOR catalytic site. After this traverse, mSin1 engages in extensive interactions with mLST8, beginning with a short complementary β-strand, followed by an extended section, which wraps approximately ¾ of the way around the peripheral surface of mLST8, and finally the whole visible part of mSin1 ends in a short α-helix before merging into the non-visualized CRIM domain (Figure 3A).

**Figure 3.**
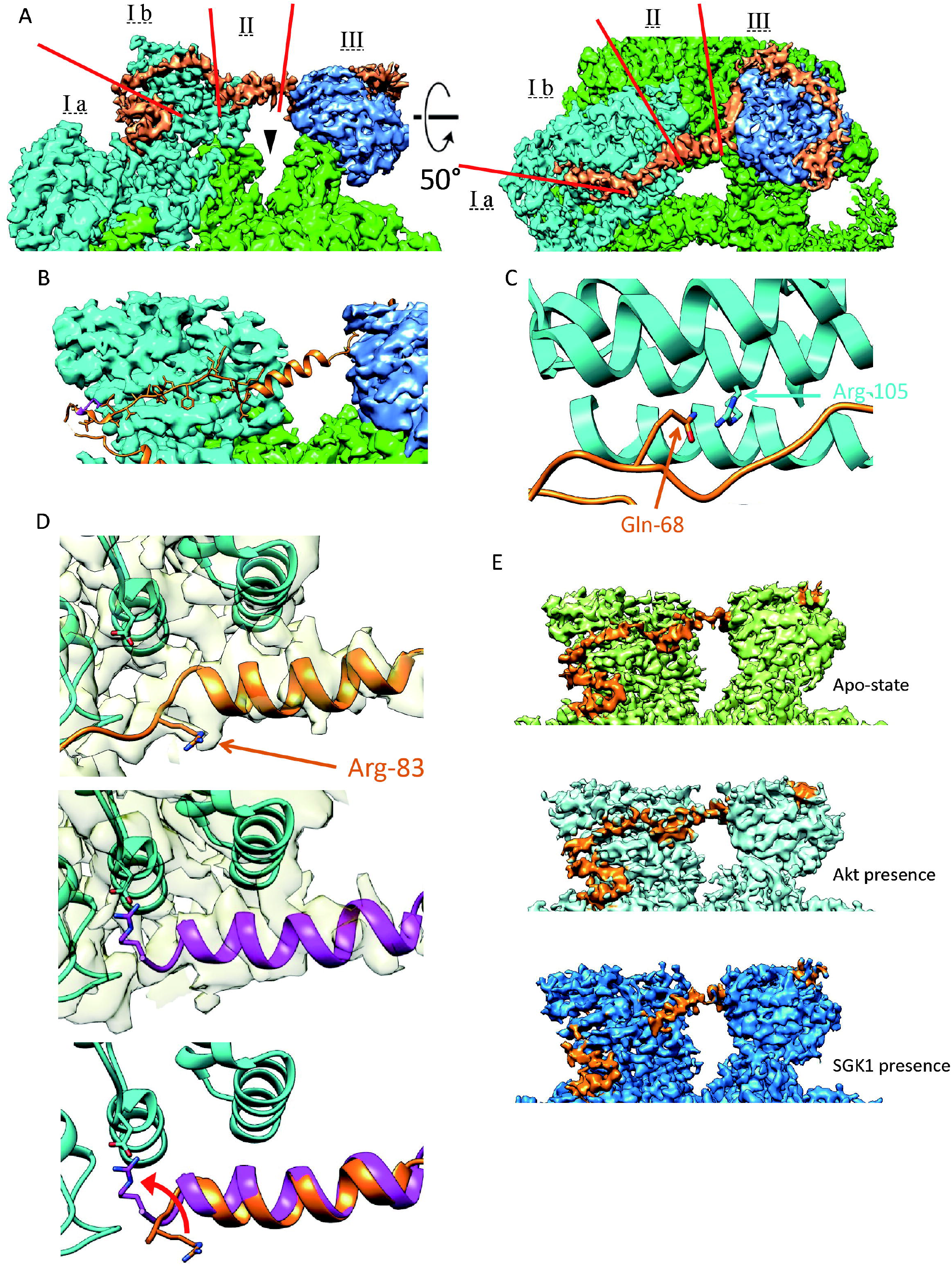
Structural details of the N-terminal domain of mSin1 (orange): (A) The N-terminus of mSin1 is shown in two views, comprising a Rictor-interacting section (I), a bridge section connecting Rictor and mLST8 (II) and an mLST8-interacting section (III). The Rictor interacting section is divided into a pair of short α-helices (I a) and an extended “string” domain (I b, see text for details). The kinase cleft of mTOR is shown by arrowhead. (B) String domain stays at the periphery of Rictor HR1, shown in the atomic model. (C) mSin1/Gln-68, as a key amino acid to mediate SGK1 interaction with and phosphorylation by mTORC2, points toward Rictor/Arg-105 in the apo-state. (D) The mSin1 Arg-83 side chain and nearby backbone with density map semi-transparent in the background are shown (upper and middle panels); nearby Rictor sequences are also seen (colored in cyan), including side chain of Asp-1679. Note marked difference in position of Arg-83 side chain in the absence (top panel) and presence (middle panel) of SGK1. mSin1 is modeled in orange in the apo-complex and purple in the SGK1 co-complex. Lower panel shows a superposition of the apo- and co-complexes (with density map hidden for clarity), emphasizing the SGK1-induced rotation in Arg-83 and formation of a salt bridge with Rictor/Asp-1679. (E) mSin1 string domain is observed in the apo-complex (top panel) and co-complex with Akt (middle panel) of mTORC2, but becomes unobservable in the co-complex with SGK1 (bottom).

### The mSin1 N-terminal domain changes conformation in the presence of SGK1 but not Akt and a new side chain interaction with Rictor is observed

In the co-complexes with Akt and SGK1, the substrates themselves were not resolved, likely due to movement and flexibility of the substrate binding domains themselves or as a consequence of freezing. Biochemical data demonstrate the physical interaction of the substrates with mTORC2. (Figure S3). The overall volume and shape of the complexes were similar to the apo-complex (Figures S4 and S5), however, the detailed relationship of the mSin1 N-terminal domain with Rictor was markedly altered in the SGK1 co-complex. First, the string domain upstream of Arg-83 including Gln-68 becomes unobservable, likely due to increased flexibility. Akt, in contrast, has no such effects: the string domain in its presence looks very similar to the apo-complex (Figure 3E, second panel). Second, the Arg-83 side chain shows a large rotation toward a negatively charged patch of Rictor HR3, coming into close proximity to Asp-1679, with which it appears to form a salt bridge (Figure 3D, middle and lower panel). In order to explore the functional role of Arg-83 in differential regulation of SGK1 and Akt, we tested the ability of mSin1/Arg-83-Ala, to support mTORC2 kinase activity by using mSin1-deficient HEK-293T cells (Gleason et al., 2019). These cells are defective in phosphorylation of both SGK1 and Akt, which is restored by WT mSin1 (Gleason et al., 2019) (Figure 4). Consistent with the specific role of mSin1/Arg-83 in SGK1 phosphorylation, Akt but not SGK1 phosphorylation was restored by mSin1/Arg-83-Ala. Together with prior literature (Lu et al., 2011b; Lu et al., 2010), these data support a model in which SGK1 binding to the string domain disrupts several weak interactions with Rictor (including mSin1/Gln-68 with Rictor/Arg-105), while concomitantly inducing a conformational change that results in a new stabilizing interaction.

**Figure 4.**
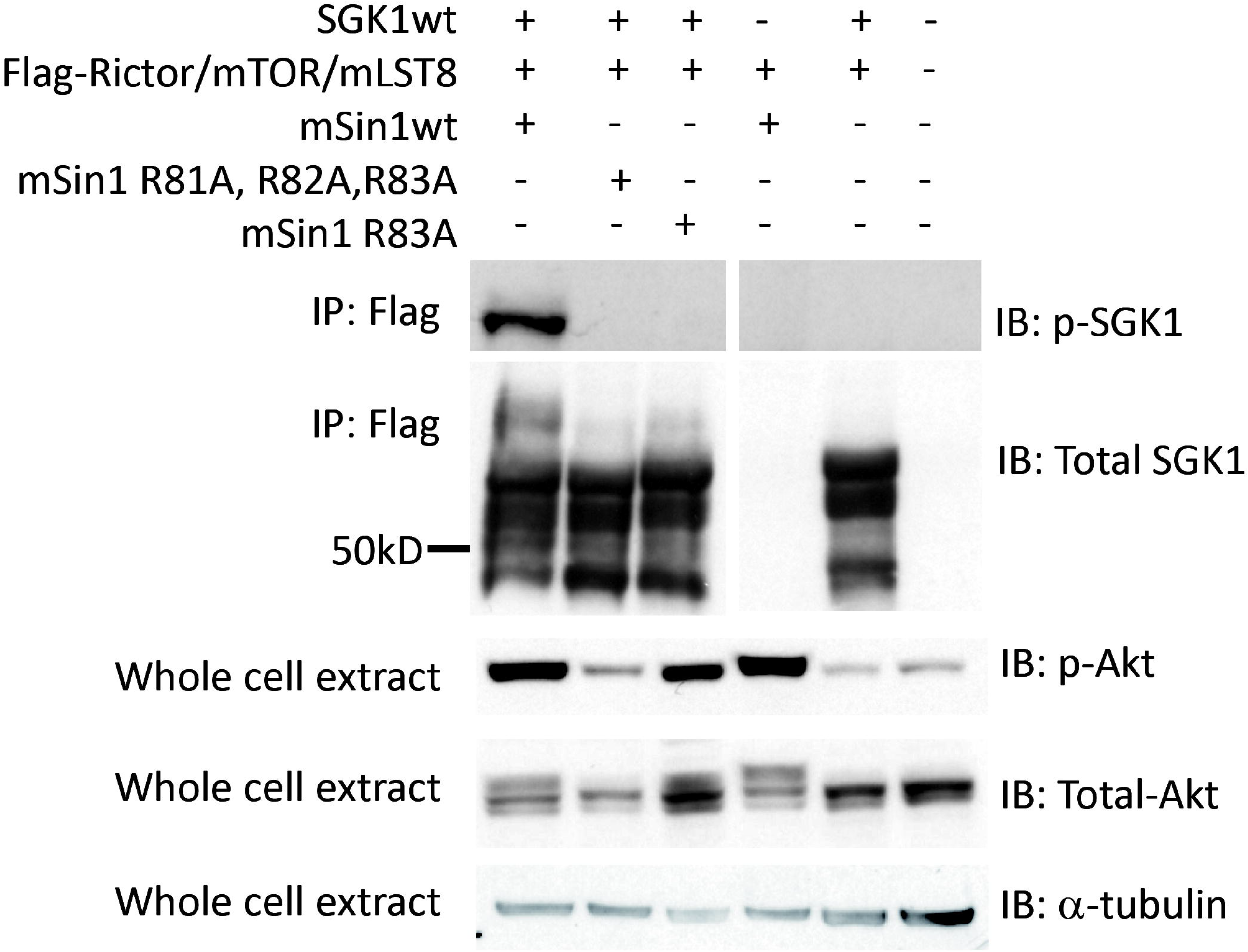
mSin1/Arg-83 is required for SGK1 but not Akt phosphorylation. mSin1 knock-out HEK293T cells were cotransfected with Myc-tagged Rictor, SpyCatcher003-tagged mTOR, HA-tagged mLST8, HA-tagged mSin1 (wild-type, R81A/R82A/R83A triple mutant and R83A single mutant), and FLAG-tagged SGK1. Flag-Immunoprecipitation and whole cell extracts were prepared and subjected to immunoprecipitation with anti-FLAG antibody followed by immunoblotting (IB) as shown, and further described in “Experimental Procedures”. Blots were probed with antibodies against phosphorylated SGK1, total SGK1, phosphorylated Akt, total Akt, and α-tubulin, respectively.

## Discussion

mTORC1 and 2 behave as distinct signaling kinases despite sharing a common core catalytic subunit, mTOR. Previous work on the mTORC1 structure has provided a detailed picture of key specificity determinants with high resolution, using a combination of X-ray crystallography and cryo-EM (Aylett et al., 2016; Yang et al., 2013). A previous structure of mTORC2 using cryo-EM reached a resolution of 4.9 Å (Chen et al., 2018), and a paper that was published while this manuscript was in preparation reached a resolution of 3.0 Å (Scaiola et al., 2020). Our present reconstruction confirms key features identified in those structures, and reveals structural features not previously visualized, providing new mechanistic insight. Instrumental in obtaining our more comprehensive density map was the use of a recently developed on-grid capture purification system utilizing the self ligating “Spycatcher-Spytag system”. To our knowledge, this is the first adaptation of this technology to purifying a large protein complex (∼1.2 MDa) for cryo-EM study, as described in detail in Methods and Results. The approach significantly conocentrates the sample on the grid, minimizes non-specific binding, and protects the delicate mTORC2 particles from both the air-water interface and the graphene oxide surface. The net effect is to enhance the reconstruction resolution and to improve the density quality of peripheral parts of the complex.

In the overall structure, we show that mTORC2 is not a rigid structure, but rather, using 3D variability analysis, we visualized the two halves moving in dynamic “breathing” modes. This movement likely represents a correlate of the dynamic flexibility of mTORC2, and may contribute to the differential behavior of mTORC1 and mTORC2. It seems likely to play a role in limiting the resolution of the overall structure of mTORC2 in the present and earlier structures (Chen et al., 2018; Scaiola et al., 2020). Further studies are required to investigate the biological significance of these movements, which may answer some remaining questions such as whether the interaction patterns of the two mTOR monomers differ between mTORC1 and mTORC2 (Chen et al., 2018). Movement of the mSin1 N-terminal region is associated with its propensity to become increasingly flexible in response to one of its substrates, SGK1, as addressed further below.

Our structure provides novel insight into the atomic details of the interaction between Rictor and the mTOR FRB domain, which is critical both for the relative rapamycin insensitivity of mTORC2, and for its recruitment of S6-kinase and 4EBP1. In particular, specific contacts were observed between Ser-1624 and Ser-1625 of Rictor and mTOR/Thr-2098 (Figure 2D), which preclude interaction with FKBP12-rapamycin or mTORC1 substrates (Yang et al., 2017). We also confirmed close apposition of Rictor/Val-1627 and mTOR/Ser-2035 that likely blocks access as well (Figure 2D). Thus, Rictor plays a central role in mTORC2 specificity by limiting access to mTORC1 substrates and both endogenous and pharmaceutical small molecule ligands (Loewith et al., 2002; Sarbassov et al., 2004).

While Rictor plays a central role in blocking access of mTORC1 substrates, mSin1 appears to be the principal mTORC2 component mediating substrate recruitment. The CRIM domain of mSin1 has been shown to interact with all three major mTORC2 substrates (Cameron et al., 2011b; Tatebe et al., 2017), and we showed previously that the N-terminal region interacts selectively with SGK1 (Lu et al., 2011a). Our structure now reveals for the first time the N-terminal extended strand or “string” (SGK-Targeting, Rictor InteractING) domain of mSin1 (aa 66-86), which includes Gln-68 shown previously to be required specifically for activation of SGK1, but not Akt or PKC (Lu et al., 2011a). Here we show that in the apo-complex, the Gln-68 side chain reaches toward Rictor/Arg-105, consistent with a stabilizing interaction. Additionally, we observe multiple side chains between Gln-68 and Thr-86 forming weak interactions with Rictor HR1. Thr-86, the side chain of which interacts with Rictor/Glu-1675, constitutes the beginning of the α-helical bridge segment, and provides an important anchor for the bridge, which traverses the deep catalytic cleft toward mLST8. It is notable in this context that phosphorylation of Thr-86 has been suggested to impair mTORC2 integrity (Liu et al., 2013). This issue is, however, controversial as others have found that phosphorylation of Thr-86 enhances mTORC2 activity (Yang et al., 2015). The structural correlates of this apparently context-dependent effect of Thr-86 phosphorylation are unclear at this time, as Thr-86 was unphosphorylated in our structures and those of others (Scaiola et al., 2020).

Comparison of apo-complex and co-complex structures reveals a marked effect of SGK1 on mTORC2 conformation in the string domain: Arg-83, which is quite far from any potential interacting residue in the apo-complex, undergoes a large rotation toward a negative patch in HR1, which includes Rictor/Asp-1679, with which it appears to form a salt bridge. Concomitant with this conformational change, the string domain upstream of Arg-83 becomes invisible, likely due to increased flexibility as it interacts with SGK1 and comes free from its moorings to Rictor HR1. We propose that the new Arg-83-Asp-1679 interaction replaces other (weaker) string domain interactions to maintain the stability of mTORC2, and further suggest that the substrate-induced increase in string domain flexibility allows it to coordinate with the CRIM domain in binding SGK1 and present the hydrophobic motif to the mTOR catalytic cleft. The functional importance of Arg-83 is further supported by the observation that an Ala mutant of that residue selectively disrupts SGK1 but not Akt phosphorylation.

It is of interest to consider the implications of our structural observations for mTORC2 activation of its distinct substrates. Akt, SGK and conventional PKCs are all mTORC2 targets, however, there are significant differences in the subcellular location of activation, relevant upstream effectors and physiological activating conditions (Fu and Hall, 2020; Gleason et al., 2019). Indeed, recent data have identified hormonal and non-hormonal effectors that selectively stimulate mTORC2 phosphorylation of SGK1 but not Akt (Gleason et al., 2019; Sorensen et al., 2019). Akt regulation is substantially controlled by its N-terminal PH domain, which binds to phosphatidylinositol-3,4,5-triphosphate (PIP3), mediates colocalization with mTORC2 at the plasma membrane, and together with CRIM domain interaction brings kinase and substrate together (Cameron et al., 2011a; Pearce et al., 2010). SGK1 has a distinct N-terminal domain, which binds monophosphorylated phosphoinositides but not PIP3 (Pao et al., 2007). Interaction with both the mSin1 string and CRIM domains is essential for its activation (Cameron et al., 2011b; Lu et al., 2011a). It seems likely that string domain binding and the ensuing conformation change in mSin1 play an important role in selective hormonal regulation of SGK1 (for example by angiotensin II) (Gleason et al., 2019), and responses to internal physiological stressors such as severe cold (Allu et al., 2021) or rapid changes in electrolytes (Sorensen et al., 2019). Further studies will be needed to fully visualize the detailed interactions of SGK1 with mSin1 that support its selective regulation by mTORC2.

## Methods

### Reagents

Antibodies: anti-Rictor (2114, CST), anti-mTOR (2972, CST), anti-HA (3724, CST), anti-Akt (4691, CST), anti-phospho-Ser473 Akt (4060, CST), anti-phospho-Ser422 SGK1 (SC-16745-R, SCBT), horseradish peroxidase (HRP)-labeled anti-rabbit secondary antibodies (7074, CST).

Other reagents: anti-FLAG M2 agarose affinity resin (A2220, Sigma), 3×FLAG peptide (F4799, Sigma), Expi29™ expression medium (A1435101, Thermo Fisher Scientific), ExpiFectamine™ 293 transfection kit (A14525, Thermo Fisher Scientific), Opti-MEM™ I reduced serum medium (31985062, Thermo Fisher Scientific), dulbecco’s modified eagle medium (DMEM) with high glucose (4.5 g/Liter) (CCFAA005, UCSF Cell Culture Facility), fetal bovine serum (FBS; 11650, Atlanta Biologicals), penicillin-streptomycin (30-002-CI, Corning), L-glutamine (25-005-CI, Corning), trypsin (25-052-CI, Corning), polyethylenimine (PEI; 24765, Polysciences), complete EDTA-free protease inhibitor cocktail (11697498001, Roche), PhosSTOP phosphatase inhibitor (04906837001, Roche), Bradford assay reagent (1856209, Thermo Fisher Scientific), ECL western blotting detection reagent (RPN2106, GE Heathcare), Lambda protein phosphatase (P0753, NEB), AZD8055 (S1555, Selleckchem), Superose 6 increase 3.2/300 GL (29091598, GE Healthcare) and Superdex 200 increase 3.2/300 GL (28990946, GE Healthcare).

### Protein expression and purification

To produce human mTORC2 protein, plasmids pcDNA3-FLAG-mTOR, pRK5-myc-Rictor, pcDNA3-mSin1(1.1)-HA, pRK5-HA-mLST8 and pDEST14-SpyCatcher003 were gifts from Jie Chen (Addgene plasmid #26603), David Sabatini (Addgene plasmid #11367), Jie Chen & Taekjip Ha (Addgene plasmid #73388), David Sabatini (Addgene plasmid #1865), and Mark Howarth (Addgene plasmid #133447), respectively. Site-directed mutagenesis was performed on pRK5-myc-Rictor to replace its original tag and generate pRK5-FLAG-Rictor. The ORF of SpyCatcher003 was subcloned from pDEST14-SpyCatcher003 into pcDNA3-FLAG-mTOR to replace its original tag and generate pcDNA3-SpyCatcher003-mTOR plasmid. All DNA constructs were verified by DNA sequencing.

Expi293F™ cells (A14527, Thermo Fisher Scientific) were grown in Expi29™ expression medium in sterile polycarbonate Erlenmeyer flasks (89095, VWR) in a shaker (MaxQ 416 HP, Thermo Scientific Scientific) at 120 rpm at 37°C and 8% CO_2_. The four plasmids, pcDNA3-SpyCatcheer003-mTOR, pRK5-FLAG-Rictor, pcDNA3-mSin1(1.1)-HA and pRK5-HA-mLST8, were co-transfected in 100 μg per 100 ml Expi293F™ cells using ExpiFectamine™ 293 transfection kit and Opti-MEM™ I reduced serum medium, according to manufacturer’s instructions. After cultured for 72 hours, cells were harvested and lysed with lysis buffer (0.1% CHAPS, 40 mM HEPES pH 7.5, 1 mM EDTA, 120 mM NaCl, 50 mM NaF, 10 mM Na Pyrophosphate, 10 mM Glycerophosphate) containing complete protease inhibitor cocktail at 4°C for 30 minutes. After centrifugation at 17,000×g, 4°C for 10 minutes, the supernatant was collected and added to anti-FLAG M2 agarose affinity resin. After incubation with rolling for 2 hour at 4°C, the resin was thoroughly washed with wash buffer (30 mM HEPES pH 7.4, 0.5 mM EDTA, 200 mM NaCl) and the protein complex was eluted from the resin with 0.25 mg/mL 3×FLAG peptide in elution buffer (30 mM HEPES pH 7.4, 0.5 mM EDTA, 200 mM NaCl, 1.0 mM TCEP) for 30 minutes at 4°C. The protein was concentrated to 8.0 mg/ml using 100k molecular weight cutoff Amicon columns (UFC510024, Millipore) and further purified by gel filtration chromatography (Superose 6 increase 3.2/300 GL) in the buffer containing 30mM HEPES pH 7.4, 200mM NaCl, and 0.5mM EDTA. The peak fractions were concentrated to 1.1 mg/ml and ready for cryo-EM studies. For kinase assay, 10% (v/v) glycerol was added to purified protein aliquots for storage at −80°C.

To produce inactive human kinase dead Akt1 and wild type SGK1 protein, site-directed mutagenesis was conducted on pcDNA3-HA-Akt1-K179M (Addgene plasmid #73409) and pcDNA3-SGK1-V5-6xHis to change their original tags into FLAG tag. 293T cells were grown in DMEM supplemented with 10% FBS, 1% L-glutamine and 1% penicillin-streptomycin at 37°C, 5% CO_2_ and transfected with the plasmid pcDNA3-FLAG-Akt1-K179M was transfected into 293T cells using PEI. Expi293F™ cells were cultured and transfected with the plasmid pcDNA3-SGK1-FLAG as described above. Both cells were harvested and lysed in lysis buffer (0.1% CHAPS, 40 mM HEPES pH 7.5, 1 mM EDTA, 120 mM NaCl, 50 mM NaF, 10 mM Na Pyrophosphate, 10 mM Glycerophosphate) containing complete protease inhibitor cocktail at 4°C for 30 minutes. The clarified cell lysates were incubated with anti-FLAG M2 agarose affinity resin for 2 hours at 4°C and eluted with 0.25 mg/mL 3×FLAG peptide in elution buffer (30 mM HEPES pH 7.4, 0.5 mM EDTA, 200 mM NaCl, 1.0 mM TCEP) for 30 minutes at 4°C. Akt1 and SGK1 were concentrated to 4.5 mg/ml and 3.5 mg/ml respectively, using 30k molecular weight cutoff Amicon columns (UFC503024, Millipore). To dephosphorylate them, proteins were incubated with lambda phosphatase for 1 hour at room temperature and overnight at 4°C. Dephosphorylated Akt1 and SGK1 were further purified by gel filtration chromatography (Superdex 200 increase 3.2/300 GL) in the buffer containing 30mM HEPES pH 7.4 and 200mM NaCl. The peak fractions were concentrated to 1.7 mg/ml for Akt1 and 0.52 mg/ml for SGK1, directly forwarded to co-complex construction with mTORC2, or added with 10% (v/v) glycerol and stored at −80°C for kinase assay.

To get purified mTORC2+Akt co-complex and mTORC2+SGK1 co-complex, the raw mTORC2 protein after 3×FLAG peptide elution and concentration, ∼8.0 mg/ml, was incubated with Akt or SGK1 purified above at molar ratio of 1:2. After incubation for 2 hours at 4°C, the mixture was further purified by gel filtration chromatography (Superose 6 increase 3.2/300 GL) in the buffer containing 30mM HEPES pH 7.4, 200mM NaCl, and 0.5mM EDTA. The peak fractions of expected molecular weight were checked by SDS-PAGE and Coomassie blue staining, and fractions containing the correct co-complex were concentrated to 0.5 mg/ml and ready for cryo-EM studies.

### In vitro phosphorylation assay

The in vitro kinase assays were performed in the reaction buffer containing 25 mM HEPES (pH 7.4), 100 mM Potassium Acetate, and 2 mM MgCl_2_. In a 45 μl reaction system, 0.55 μg purified mTORC2 was mixed with 0.5 ug purified Akt1 (K179M) or SGK1, and added with 10 μM AZD8055 where indicated. Reactions were initiated by adding 0.5 mM ATP and incubated for 30 min at 37°C. Reactions were stopped by adding Laemmli sample buffer and followed by SDS-PAGE and immunoblotting.

### Immunoprecipitation

Knock-out of mSin1 HEK293T cell was described before [ref Gleason 2019 JCS]. mSin1 knock-out HEK293T cells were transfected by the plasmids in this study by PEI following the provider’s instruction. Transfected cells were grown for protein expression then lysed in binding buffer (50 mM Tris-HCl, pH 7.5, 10% glycerol, 1 mM EDTA, 2 mM DTT, 150 mM NaCl, and 0.1% CHAPS) for 30 min. After centrifugation, the supernatants were collected and incubated with the anti-FLAG M2 agarose affinity resin. The immunoprecipitates were collected by centrifugation, washed three times, and boiled for 5 min in 50 μl of cracking buffer (50 mM Tris-HCl, pH 7.0, 10% glycerol, 2% SDS, 2% β-mercaptoethanol). Immunoblotting was carried out by separating the immunoprecipitates on 10% polyacrylamide gels as described using a Bio-Rad Mini-Gel apparatus and transferring them electrophoretically to Hybond-C Extra membranes (GE Healthcare) using a Trans-Blot apparatus (Bio-Rad). The membranes were incubated to block nonspecific binding in 5% nonfat dry milk in T-PBS (1.5 mM KH_2_PO_4_, 8 mM Na_2_HPO_4_, 2.7 mM KCl, 130 mM NaCl, and 0.1% Tween 20) with gentle agitation for 1 h at room temperature and probed by Western blotting (for endogenous and transfected proteins, as described in the figure legends), using antibodies against SGK1 (Sigma), phospho-SGK1 (S422) (Santa Cruz Biotechnology), Akt, phospho-Akt, mSin1 (Bethyl Laboratories). After washing with T-PBS, the membranes were incubated with peroxidase-conjugated goat anti-rabbit IgG in T-PBS for 1 h, washed three times in T-PBS, and incubated with the ECL western blotting detection reagents according to the manufacturer’s instructions.

### Immunoblotting

The protein samples were subjected to Laemmli sample buffer, run on SDS-PAGE, and transferred to a PVDF membrane pretreated by methanol for activation. After blocked with 5% non-fat milk in TBST for 30 minutes at room temperature, the membrane was probed with primary antibodies against HA tag, mTOR, Rictor, Akt, or phospho-Ser473 Akt (1:1000 dilution) overnight at 4°C followed by HRP-conjugated anti-rabbit secondary antibodies (1:5000 dilution) for 1 hour at room temperature. The protein bands were treated by ECL western blotting detection reagents and visualized by ChemiDoc imaging system (Bio-Rad). Densitometry of the band intensity was further determined via Image J software.

### Electron-microscopy data acquisition

For negative staining, 2.5 µl of the purified mTORC2 was applied to glow-discharged EM grids covered by a thin layer of continuous carbon film (TED Pella, Inc.) and stained with 0.75% (w/v) uranyl formate solution as described. EM grids were imaged on a Tecnai T12 microscope (Thermo Fisher Scientific) operated at 120 kV with a camera UltraScan4000 (Gatan Inc.). Images were recorded at a magnification of ×52,000, in a 2.23-Å pixel size on the specimen. Defocus was set to −1.5 µm. For cryo-EM, 3 µl of purified mTORC2 sample (∼0.8 mg/ml) was applied to Spytag003 immobilized affinity grids, which were fabricated as described previously (Wang et al., 2020). The grids were dip washed twice followed by adding 3 µl buffer onto grid to prevent drying. They were blotted by Whatman No. 1 filter paper and plunge-frozen in liquid ethane using a Mark IV Vitrobot (Thermo Fisher Scientific) with blotting times of 3–6 s at room temperature and over 90% humidity. Cryo-EM datasets were collected using SerialEM and EPU in different microscopes all equipped with Field Emission source and K3 camera (Gatan Inc.) operated by SLAC national facility.

### Image processing and model building

For all cryo-EM datasets, movies were motion-corrected by MotionCor237. Motion-corrected sums without dose weighting were used for defocus estimation by using CTFFIND4, incorporated in the cisTEM. Motion-corrected sums with dose weighting were used for all other image processing. All datasets were processed similarly. In summary, particles were picked automatically, refinement package created and 2D class averages, initial model generation and autorefinement by following the workflow in cisTEM (Grant et al., 2018; Zivanov et al., 2018). Final maps were refined, reconstructed and sharpened in cisTEM. Resolution was estimated by FSC = 0.143 criterion. Atomic model of mTORC2 (PDB: 5ZCS) was used as a starting model followed by manually adjusting in COOT to fit the density map, followed by iterative refinement with Phenix. real_space_refine and manually adjustment in COOT. This process was repeated until Ramachandran validation was satisfied. UCSF Chimera and Pymol were used to prepare images.

### 3D variability analysis

We carried out 3D variability analysis in cryoSPARC version 2 upon all particles used for the final 3D reconstruction from each dataset as input (Punjani and Fleet, 2021). Analytical computations were performed for the global structures. In each case, multiple modes of variability were solved, and represented as conformational changes. To visualize the transformation of density, we calculated twenty reconstructions with a filter resolution of 6 Å. A movie that combines these reconstructions as frames was generated in Chimera for each dimension of motion.

## Acknowledgements

Some of this work was performed at the Stanford-SLAC Cryo-EM Center (S^2^C^2^) supported by the NIH Common Fund Transformative High Resolution Cryo-Electron Microscopy program (U24 GM129541). We thank David P. Bulkley, Glenn Gilbert and Matthew Harrington from UCSF Keck Foundation Advanced Microscopy Laboratory for their assistance. We also thank Louella Lee for administrative support. This work is supported by the UCSF cryoEM facility grants: 1S10OD026881, 1S10OD020054, 1S10OD021741; NIH grants R01-DK56695 (D.P.), R35GM118099 (D.A.A.), R35GM140847 (Y.C); Grant from the James Hilton Manning and Emma Austin Manning Foundation (D.P.); Y.C. is an Investigator of Howard Hughes Medical Institute.

## Figure Legends

Figure S1

Purification and characterization of mTORC2 (A) The flow-chart of purification procedure. (B) The size-exclusion gel filtration Superose 6 3.2/300 profile of mTORC2 after flag pull-down. The void and sample peaks are indicated. (C) Gel filtration fractions were resolved by the SDS-PAGE 4-15% gradient gel and further stained by Coomassie blue. The desired fractions were selected for on-grid binding and cryo-EM study. (D) Kinase assay of mTORC2. Purified human mTORC2 were used as kinase. Human Akt and SGK1 were used as substrates. AZD8055 was applied as indicated to confirm the mTOR specific kinase activity. The phosphorylation of Akt and SGK1 was detected by immunoblotting using antibody against Akt, phospho-Ser473 Akt, SGK1 and phospho-Ser422 SGK1, respectively. The integrity of mTORC2 was verified by immunoblotting. (E) The atomic model of mTORC2 in the apo-state according to the high-resolution density map. Subunits are colored as indicated.

Figure S2

The cryo-EM data of mTORC2 in the apo-state. Upper left panel: A typical cryo-electron micrograph (scale bar 63 nm) of frozen hydrated mTORC2. Inserted below are representative 2D class averages. Upper right panel: Fourier power spectrum calculated from the micrograph matched with simulated Thon rings. Lower left panel: dFSC from different Fourier cones. Each curve indicates a different direction. dFSC of mTORC2 in the apo-state is presented. Lower right panel: the density map of final reconstruction of mTORC2 in the apo-state.

Figure S3

mTORC2 and its substrates interact with each other to form co-complexes: (A) Purified mTORC2 and Akt were incubated together and analyzed by size exclusion Superose 6 chromatography (left panel). The fractions according to the peak shown up in the FPLC profile were resolved by SDS-PAGE 4-15% gradient gel and stained by Coomassie blue prior to use for cryo-EM (Middle panel). Fractions used for cryo-EM analysis were immunoblotted by antibodies as indicated to verify the subunits of mTORC2 and Akt respectively (Right panel). (B) Purified mTORC2 and SGK1 were incubated together and analyzed by size exclusion Superose 6 chromatography (left panel). The fractions according to the peak shown up in the FPLC profile were resolved by SDS-PAGE 4-15% gradient gel and stained by Coomassie blue prior to use for cryo-EM (Middle panel). Fractions used for cryo-EM analysis were immunoblotted by antibodies as indicated to verify the subunits of mTORC2 and SGK1 respectively (Right panel).

Figure S4

Cryo-EM data of mTORC2 in the presence of SGK1. Upper left panel: A typical cryo-electron micrograph (scale bar 63 nm) of frozen hydrated co-complex of mTORC2 with SGK1. Upper right panel: Fourier power spectrum calculated from the micrograph matched with simulated Thon rings. Lower left panel: dFSC from different Fourier cones. Each curve indicates a different direction. dFSC of co-complex with SGK1 is presented. Lower right panel: the density map of final reconstruction of co-complex with SGK1.

Figure S5

Cryo-EM data of mTORC2 in the presence of Akt. Upper left panel: A typical cryo-electron micrograph (scale bar 63 nm) of frozen hydrated co-complex of mTORC2 with Akt. Upper right panel: Fourier power spectrum calculated from the micrograph matched with simulated Thon rings. Lower left panel: dFSC from different Fourier cones. Each curve indicates a different direction. dFSC of co-complex with Akt is presented. Lower right panel: the density map of final reconstruction of co-complex with Akt.

Figure S6

Raptor does not block rapamycin-FKBP12 binding to the mTOR FRB domain. Top panel: Three views of mTOR FRB domain colored in red and Raptor colored in dark yellow in the absence of rapamycin. Note that mTOR/Thr-2098 (in FRB domain) is exposed, unlike in the presence of Rictor. Lower panel: Three views of mTOR FRB domain bound by rapamycin-FKBP12 complex (purple) in the presence of Raptor.

